# Minimizing time in culture: A prototypic autologous manufacturing workflow for monoclonal iPSC lines within seven weeks

**DOI:** 10.64898/2026.08.04.741960

**Authors:** Diana Haberhausen, Christian Wöhle, Constanze Raab, Carsten Ludwig, Tristan Kuchler, Sofie Barth, Ullrich Wüllner, Andreas Bosio, Hannah Johannsen, Sebastian Knöbel

## Abstract

Induced pluripotent stem cells (iPSCs) hold great promise for both allogeneic and autologous cellular therapies. However, broad application and clinical translation is hindered by fragmented, complex and time-intensive workflows, resulting in high manufacturing costs, poor standardization and increased risk of genomic aberrations in derived iPSCs. In this study we developed a standardizable, automatable and time-efficient process for the derivation of monoclonal iPSC lines straight from skin including a comprehensive and cascaded QC strategy. We generated monoclonal iPSC lines derived from human skin punch biopsies of ten donors (age 49-81) via mRNA-based reprogramming that subsequently underwent comprehensive and thorough characterization of phenotypic and genetic properties. The use of a combined mechanical and enzymatic fibroblast isolation protocol and a transient non-integrative reprogramming technology allowed us to obtain 78 monoclonal iPSC lines, ready for banking, molecular characterization and further differentiation within seven weeks from initial sample processing to passage four iPSC lines. The phenotypical characterization via flow cytometry-based pluripotency marker expression and 2D-directed differentiation into the three germ layers showed low intra- and inter-donor variability over all generated lines. A combination of SNP array based CNV analysis followed by whole exome sequencing proved to be the most efficient approach for assessment of genomic integrity. Proof-of-concept experiments for closed system processing revealed that a substantial part of the most error-prone and technically demanding steps can be transferred to semi-automated, closed systems. In conclusion, the described protocol allows for time-efficient, standardizable and automatable generation of high-quality monoclonal iPSC lines from human skin punch biopsies within seven weeks, thus moving the field of autologous iPSC manufacturing one step further towards cost-efficient clinical implementation.

## Introduction

Induced pluripotent stem cells (iPSCs), as a renewable source for generation of a variety of different cell types, offer the potential to revolutionize regenerative medicine (1,2). Given the rapidly expanding range of cell types that can be derived from iPSCs (3,4), the reliable generation of iPSC lines from somatic cells has become increasingly important. Especially for autologous cell therapies, there is a high demand for having a fast, robust and streamlined workflow, which includes primary donor material preparation, reprogramming, and cell characterization with particular focus on genetic safety. Traditional reprogramming methods are often limited by low efficiency, lengthy procedures, or risks of genomic integration. While retro- and lentiviral approaches achieve high efficiencies, stable integration of viral DNA can cause persistent transgene expression and insertional mutagenesis, raising safety concerns (5), and requiring higher occupational safety standards, while mRNA-based reprogramming methods eliminate the risk of vector integration and propagation. Comparative studies of standard non-integrative reprogramming methods have demonstrated that for fibroblasts, mRNA-mediated reprogramming is the most efficient reprogramming method (6), although Sendai virus transduction (7) and episomal reprogramming (8) are still used widely (9). The latter two methods necessitate screening for viral or episomal vector residuals in the resulting iPSCs, adding considerable time in culture to the workflow, whereas mRNA- reprogrammed cells are transgene-free shortly after reprogramming due to the fast degradation of exogenous mRNA. For episomal reprogramming (6), it has been reported that more than 30% of tested iPSC lines retained plasmid fragments between passages nine and eleven, whereas following Sendai virus transduction, viral persistence in iPSCs has been observed for up to ten passages (10). The prolonged persistence of reprogramming vectors impedes early quality control (QC) and the downstream use of Sendai- or episomal-derived iPSC lines by necessitating extended time in culture to achieve vector clearance, thereby increasing the risk of chromosomal aberrations (11,12). Delbany et al. reported a significantly higher mutational burden in later passages, while most of the low-passage samples were devoid of cancer-associated mutations (13), underscoring the importance of avoiding prolonged culture time.

For these scientific and economic reasons, a streamlined and time-efficient reprogramming and iPSC expansion workflow is highly desirable. Although successfully established mRNA-based approaches work for long-lived fibroblast cell lines like BJ cells (14), reprogramming success rates are often inconsistent when applied to freshly isolated donor cells (6). Moreover, commonly published protocols require an initial fibroblast cultivation period of 14-21 days, particularly when fibroblasts are obtained via outgrowth from skin biopsies, which is associated with an increased risk of contamination, too (15). Prolonged expansion poses the risk of reduced reprogramming efficiency; with increasing population doubling times and genomic alterations may accumulate (16). Alternative approaches based on mechanical and/or enzymatic tissue dissociation enable faster tissue dissociation but frequently compromise cell purity and viability (17).

Following reprogramming, bulk iPSC populations can exhibit substantial variability in differentiation capacity, genomic stability, and persistence of residual reprogramming vectors, underscoring the importance of monoclonal derivation from a single cell to obtain a genetically and functionally homogeneous cell population. This approach enhances reproducibility, facilitates characterization, and improves the reliability of downstream experimental and translational applications. However, monoclonal derivation remains technically challenging due to limited single-cell survival (18), which necessitates careful optimization of culture conditions. Consequently, while monoclonal derivation increases experimental rigor, it also adds complexity and time to the workflow, potentially limiting scalability and throughput.

Once iPSC lines are generated, selecting appropriate iPSC lines for downstream clinical applications remains challenging, as donor-derived iPSC lines vary considerably in their suitability, especially with respect to their genetic integrity. Even iPSC lines derived from the same donor may exhibit genetic aberrations of varying extent, either due to *de novo* acquisition or by process-dependent selection during reprogramming, which is technically hard to decipher. Additionally, heterogeneity between different iPSC lines can impede subsequent differentiation efficiency or cause unexpected phenotypes in iPSC-derived cells (19,20), which can affect experimental outcomes (21) and complicate the transferability and predictability of data. Taken together, there is a critical need for stringent quality control of newly generated iPSC lines. Both, the International Organization for Standardization (ISO, (22)) and the International Society for Stem Cell Research (ISSCR, (23)) issued recommendations for minimum characterization and reporting of human PSC lines to promote harmonization and established consensus guidelines on best practices. While the phenotypic evaluation of human PSCs including morphology, the analysis of pluripotency-associated markers (undifferentiated state) and multi-lineage differentiation potential (pluripotency) is well established, the assessment of genetic integrity and oncogenic risk profiling remains challenging due to the complexity of acquired data. Aneuploidies affecting the number of whole chromosomes can be detected with conventional karyotyping or G-banding, while smaller copy number variations (CNVs) comprising several genes or regulatory elements may also have detrimental effects on affected genes but often remain undetected by these methods. Chromosomal microarrays can reveal these CNVs, but no consensus exists regarding interpretation, exclusion criteria, or threshold values. The challenge in genome wide next-generation sequencing is the identification of truly relevant or disease-associated single nucleotide polymorphism (SNPs) and insertions/deletions (indels) against the plethora of non-pathogenic polymorphisms in the genome. Thus, iPSC QC lacks standardization and currently diverse methodologies and varying evaluation criteria across the field are in use (24).

For translational efforts, automation and closed systems have become critical components of the iPSC manufacturing process. The inherent complexity, variability, operator-dependence and labor-intensity of manual iPSC generation and expansion constrain scalability, reproducibility, and regulatory compliance. These challenges underscore the need for standardized and automated and/or closed systems capable of ensuring consistency and accelerating the transition from research-scale processes to clinical-grade production.

In sum, we present an optimized and time-efficient workflow for the generation of monoclonal iPSC lines, straight from skin. This workflow addresses current issues related to skin dissociation, low reprogramming efficiency, monoclonal derivation, prolonged processing times, procedural complexity, and line-to-line heterogeneity. We first established the workflow using samples from human non-diseased donors and subsequently evaluated its applicability on relevant patient material. To ensure iPSC quality, we developed a systematic pipeline integrating phenotypic assessment and rigorous genetic integrity analysis. Towards the translational perspective, we demonstrate the feasibility of transferring several error-prone and technically demanding steps into semi-automated, and/or closed processing.

## Results

### Combined mechanical and enzymatical dissociation of human skin punch biopsies robustly lead to a pure fibroblast population within seven days

To obtain fibroblast-enriched cell populations suitable for cellular reprogramming, a rapid and efficient one-week tissue dissociation and expansion protocol was established (Figure 1A). Skin punch biopsies obtained from five non-diseased donors (ND1-ND5) and five Parkinson’s Disease patients (PD1-PD5; Table 1) were subjected to combined enzymatic and mechanical dissociation. Following dissociation overnight, cell viability remained high across samples, ranging from 63.51% to 97.63% (Figure 1B). CD90 was used as a marker associated with fibroblast identity within the cell population. The yield of CD90+ cells, varied after dissociation ranging from 978 to 18,800 cells/mm² of skin punch biopsy (Figure 1C). Specifically, this corresponded to a minimum yield of 759,330 CD90+ cells following tissue dissociation from 6-mm skin punch biopsies (ND1-ND5) and a minimum of 12,282 CD90+ cells from 4-mm skin punch biopsies (PD1-PD5). Thus, all samples provided enough CD90+ cells for reprogramming and the establishment of an early-passage fibroblast stock after a seven-day expansion phase. The proportion of CD90+ cells immediately after dissociation varied between 10.28% and 55.47% (Figure 1D). After seven days of expansion, this proportion increased to 95.44% ± 2.73% (mean ± SD) CD90+ cells (Figure 1E). Throughout the expansion phase, cells displayed fibroblast-like morphology and proliferative capacity (Figure 1F), a crucial property considering that low-proliferating or even senescent fibroblasts have limited reprogramming potential (25,26). Cells were confirmed to be free of mycoplasma contamination prior to reprogramming.

**Figure 1:**
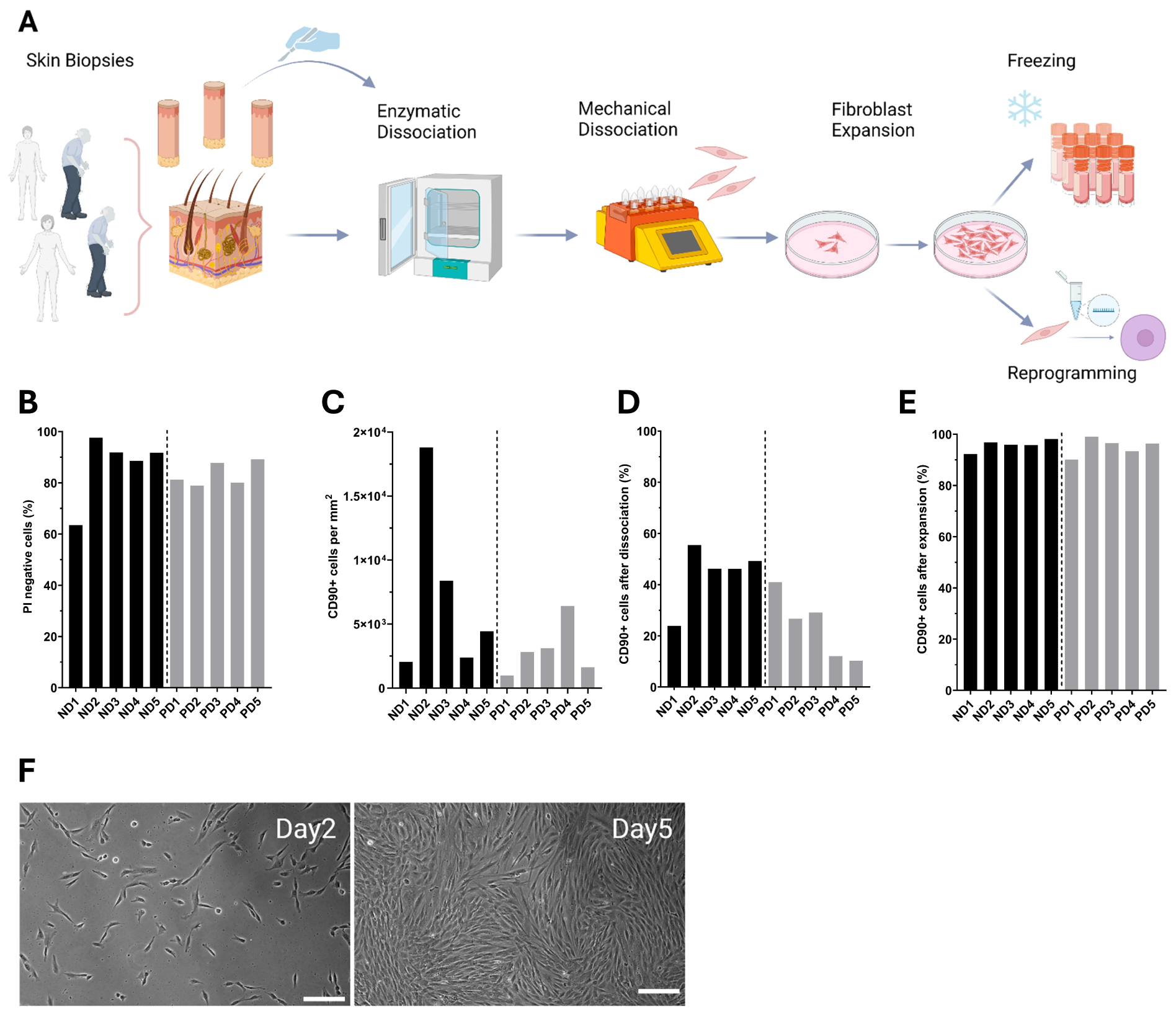
Process for skin dissociation and fibroblast expansion. (A) Schematic representation of the process for the dissociation of skin punch biopsies derived from non-diseased donors (ND1-5) and PD patients (PD1-5). (B) Viability of cells following enzymatic and mechanical dissociation. Viability was assessed by propidium iodide (PI) staining; bars represent the percentage of viable (PI-negative) cells. (C) Total number of CD90+ cells per mm^2^ skin punch biopsy after dissociation. (D) Percentage of CD90+ cells after dissociation. (E) Percentage of CD90+ cells after seven days of expansion. (F) Representative brightfield images of the fibroblast expansion. Fibroblasts two and five days after dissociation. Scale bars, 200 µm.

**Table 1:** Overview of the demographic characteristics of non-diseased donors (ND) and PD patients (PD).

| **Non-diseased donor** | **Sex** | **Age** | **Ethnicity** | **Parkinson's Disease patient** | **Sex** | **Age** | **Ethnicity** |
| --- | --- | --- | --- | --- | --- | --- | --- |
| ND1 | female | 61 | Caucasian | PD1 | male | 49 | Caucasian |
| ND2 | female | 58 | Caucasian | PD2 | male | 60 | Caucasian |
| ND3 | female | 50 | Caucasian | PD3 | male | 81 | Caucasian |
| ND4 | female | 62 | Caucasian | PD4 | male | 53 | Caucasian |
| ND5 | male | 62 | Caucasian | PD5 | male | 61 | Caucasian |

### Successful reprogramming of ten donors yields 78 monoclonal iPSC lines

Following dissociation and expansion, CD90+ cells were used as starting material for generation of monoclonal iPSC lines using mRNA-based reprogramming (Figure2A). During consecutive mRNA transfections (day0-day4), fibroblasts transitioned from an elongated to a compact, cobblestone-like morphology, indicating mesenchymal-to- epithelial transition (Figure 2C1,C2). Cells were cultured for seven additional days, leading to distinct iPSC colony formation by day eleven (Figure 2C3). Quantification of OCT3/4+ colonies on day eleven from six donors (PD1-PD5, ND5) revealed reprogramming efficiencies ranging from 6.58% to 16.02% (e.g. 6.58 colonies per 100 input cells; Figure 2D). Cyclic imaging of a representative colony further confirmed the pluripotency-associated phenotype of emerging colonies through co-staining of pluripotency- and fibroblast-associated markers. Immunofluorescence images revealed a compact iPSC colony positive for TRA-1-60, TRA-1-81, SOX2, NANOG, Ki67, and OCT3/4, surrounded by a non-reprogrammed cell layer expressing VIM, CD49a, CD90, FN, CD13 and FSP1 (Figure 2E). Reprogramming succeeded for all ten donors on the first attempt, albeit with variable percentages of TRA-1-60+ cells ranging from 3.24% to 61.57% on day eleven (27.92% ± 19.81, mean ± SD) (Figure 2G). No significant correlation was found between donor age and percentage of TRA-1-60+ cells after reprogramming (Pearson’s r = -0.20, p = 0.57; Figure 2B). Microchip-based cell sorting after reprogramming for the pluripotency-associated surface antigen TRA-1-60 successfully depleted non-reprogrammed cells while reprogrammed TRA-1-60+ cells were strongly enriched in the positive sort fraction (94.29% ± 3.68, mean ± SD; Figure 2F,H). Sorted TRA-1-60+ cells were manually seeded as single cells by limiting dilution (Figure 2C4) and were further expanded until passage four (P4), at which point the iPSC line seed stock was generated (Figure 2C5). Using this method, 78 iPSC lines were generated (Table 2).

**Figure 2:**
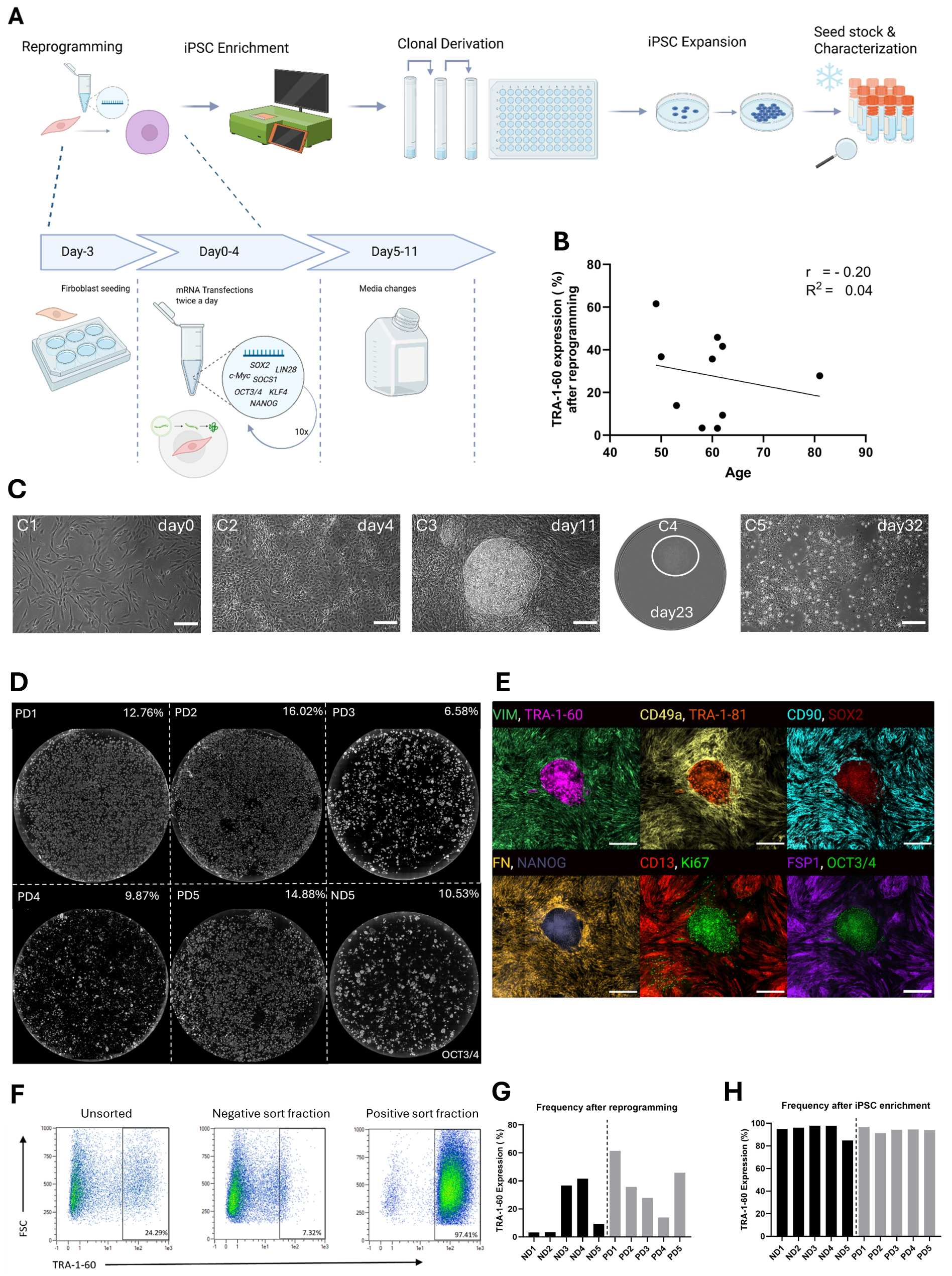
Generation of monoclonal iPSC lines. A) Schematic representation of the process for the generation of monoclonal fibroblast-derived iPSC lines. (B) Correlation between age of the donor and TRA-1-60 expression on day11 of reprogramming. Each data point represents an individual donor (n= 10; non-diseased donors and PD patients). A non-significant negative correlation was observed between donor age and TRA-1-60 expression (Pearson’s r = -0.20, p = 0.57). Linear regression line is shown with corresponding R^2^ = 0.04. (C) Representative brightfield images monitoring the reprogramming process (C1-3), monoclonal derivation (C4), and subsequent iPSC expansion (C5). (D) Overview immunocytochemistry images of a 6-well plate on day 11 of reprogramming stained for OCT3/4.Donor information and reprogramming efficiencies are shown at the top. Reprogramming efficiency (%) = (number of iPSC colonies obtained / number of starting somatic cells seeded) × 100. (E) Reprogramming of BJ fibroblasts showed a compact iPSC colony on day11 stained via cyclic immunocytochemistry imaging for pluripotency- and fibroblast-associated markers. (F) Representative flow cytometry analysis of TRA-1-60 expression in unsorted cells on day 11 of reprogramming, and in the positive and negative sort fractions following iPSC enrichment using microchip-based flow sorting. (G,H) Sort performance showing TRA-1-60 expression before (G) and after iPSC enrichment (H) for five non-diseased donors (left) and five PD patients (right) on day11 each. Scale bars, 200 µm (C); 1 cm (D); 500 µm (E).

**Table 2:**
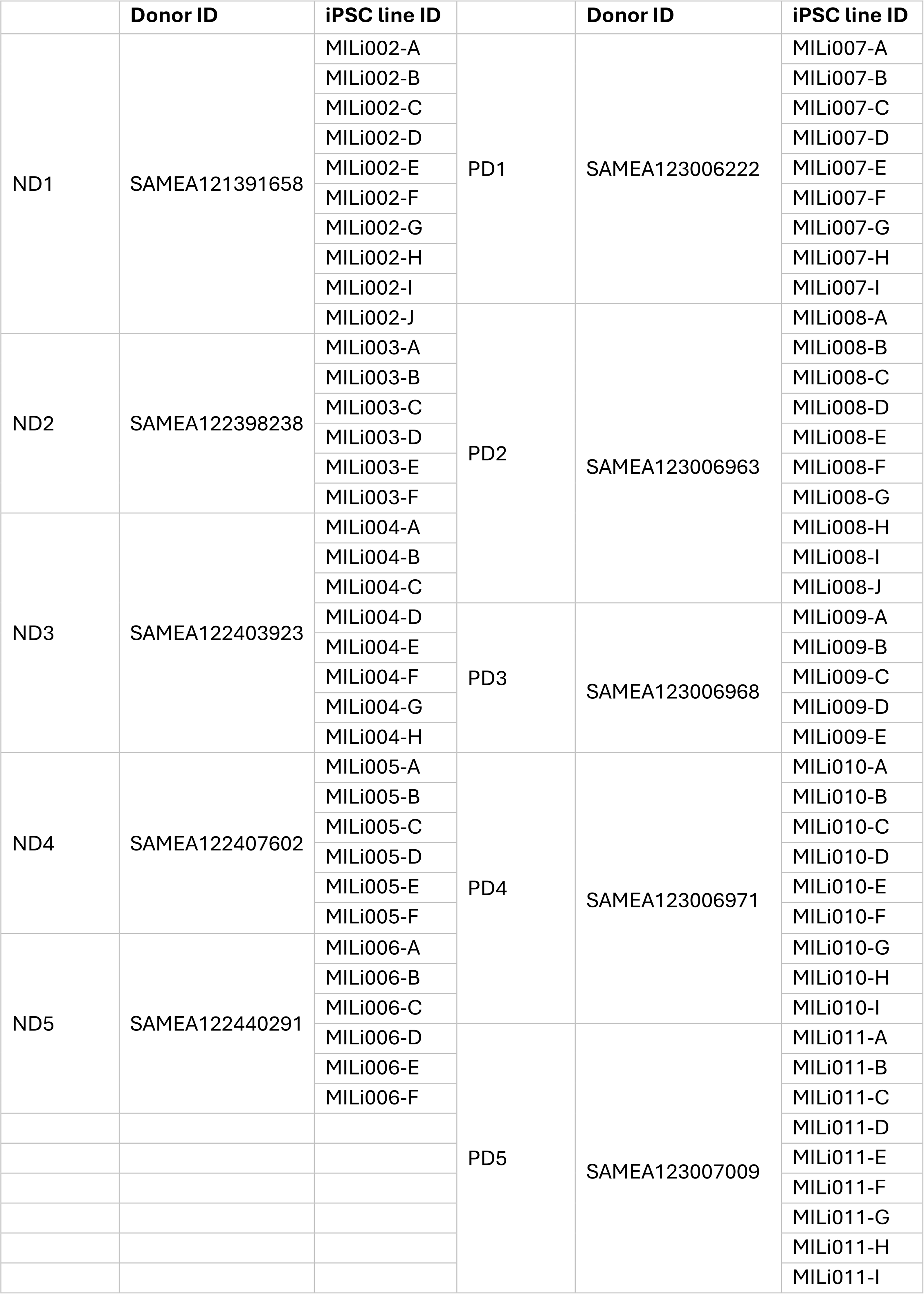
Donor and iPSC line identifier assignment for generated iPSC lines, following hPSCreg nomenclature guidelines. (**36**).

### Shortly after reprogramming, exogenous mRNAs were no longer detectable

To monitor mRNA transfection during reprogramming, GFP-mRNA was added to the mRNA cocktail as a reporter for intracellular mRNA delivery and translation. In a representative experiment, GFP expression increased progressively with consecutive transfections (Figure 3A). To initiate reprogramming, transient expression of exogenous pluripotency-associated mRNAs is required. Successful reprogramming, however, is characterized by the establishment of a self-sustaining undifferentiated state, independent of any exogenous factors. To investigate the time point when factor- independency was reached, we performed a representative RT-qPCR experiment. Exogenous mRNA levels were high during transfections (day2-4) but declined rapidly after end of transfections (Figure 3B), while endogenous pluripotency-associated gene expression increased progressively over time. By P1, following TRA-1-60 cell enrichment and a five-day cell culture, exogenous RNA was no longer detectable, while endogenous pluripotency-associated gene expression reached its maximum. While an increasing expression level could be detected during reprogramming for the endogenously transcribed genes *Lin28*, *NANOG*, *OCT3/4* and *SOX2*, no such increase was observed for *c-Myc* and *KLF4*. Notably, fibroblasts already had a basal transcription level for *KLF4* and *c-Myc* comparable to the resulting iPSCs prior to initial transfection, and the expression of both genes remained relatively unchanged throughout the reprogramming process, leading us to suspect that reprogramming might also be possible without these two factors. However, attempts to omit *c-Myc* and/or *KLF4* mRNA were unsuccessful, resulting in a lack of reprogrammed cells on day eleven of reprogramming (Fig 3C). Also, substituting *c-Myc*, with *l-Myc* remained unsuccessful in our hands (Figure 3D).

**Figure 3:**
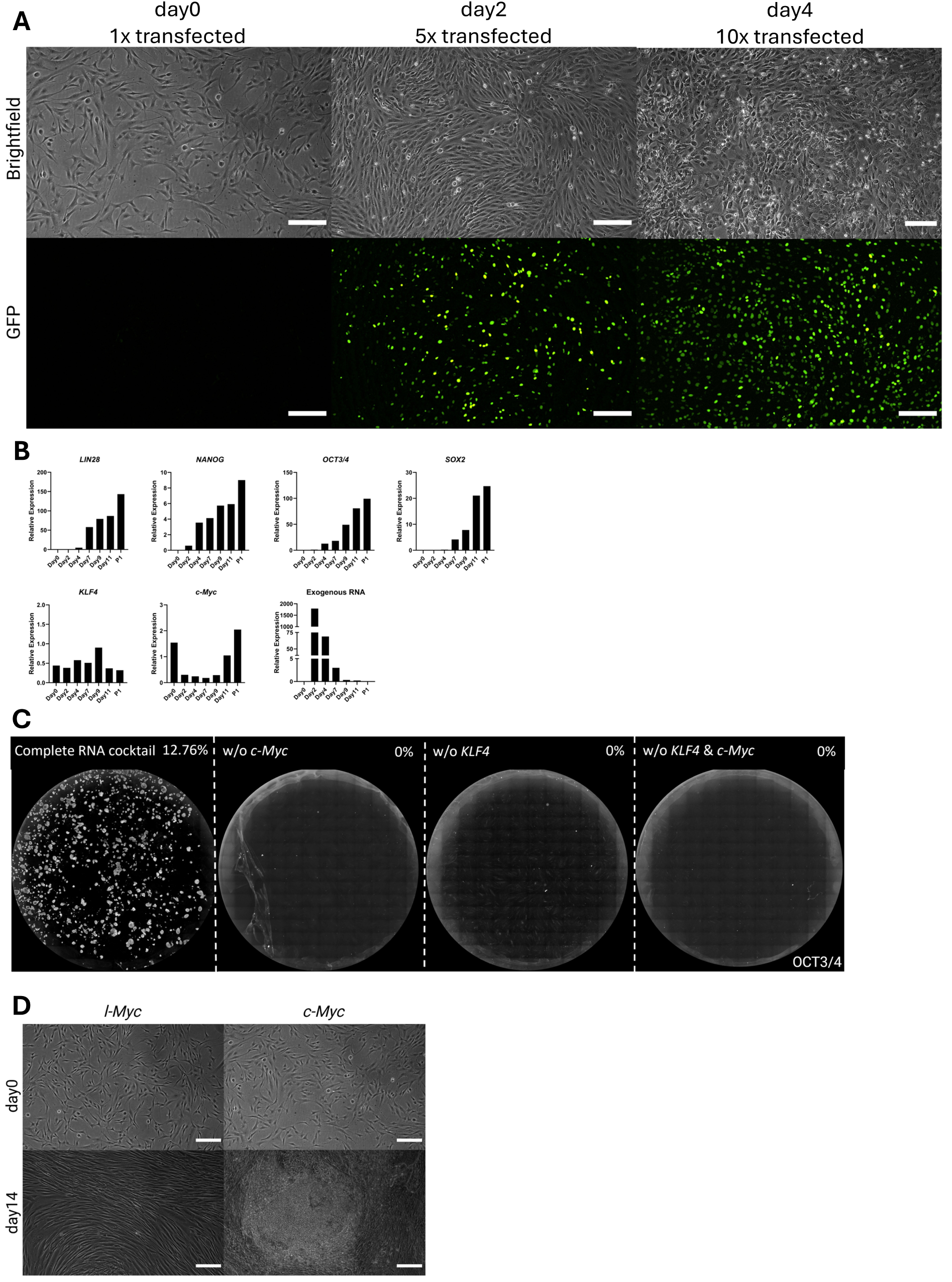
Transfection efficiency and kinetics of mRNA-mediated reprogramming. (A) Time course of GFP expression over four days following consecutive transfections with GFP-encoding mRNA. Upper panel, bright field imagines; lower panel, corresponding nuclear GFP fluorescence. (B) Relative mRNA expression of exogenous and endogenous pluripotency-associated genes during reprogramming (day0 - 11) and subsequent cell culture (P1, five days after plating of sorted cells), as measured by RT-qPCR. Results are given as relative expression over the housekeeping gene TBP (2^-ΔCT^). (C) Omission of *c-Myc* and/or *KLF4* during reprogramming. Representative immunofluorescent overview images stained for OCT3/4 of a 6-well at day11 of reprogramming with either *c-Myc*, *KLF4*, or both factors omitted from the reprogramming cocktail. Reprogramming efficiency for each condition is indicated above. (D) Attempt to replace *c-Myc* by *l-Myc*. Brightfield images of cells transfected with the complete reprogramming mRNA cocktail containing either *l-Myc* (left panel) and *c-Myc* (right panel). Scale bars, 200 µm.

### Generated iPSC lines exhibit a pluripotency-associated phenotype

Multiple iPSC lines were derived from each donor through the reprogramming procedure described above, followed by monoclonal derivation using limiting dilution and subsequent iPSC expansion. All generated iPSC lines (n=78) displayed characteristic stem cell morphology - high nuclear-to-cytoplasmic ratio, prominent nucleoli, growth in tightly packed colonies with well-defined borders and absence of spontaneously differentiated cells (Figure 4A). Flow cytometry at P4 (iPSC line seed stock) confirmed overall consistently high expression of pluripotency-associated markers SSEA-4 (99.69% ± 0.39, mean ± SD), SSEA-5 (99.55% ± 0.59), TRA-1-60 (99.38% ± 0.60), SOX2 (97.13% ± 2.73), and OCT3/4 (97.66% ± 2.78), while minimal CD15 expression (0.52% ± 0.55) indicated negligible potential for spontaneous differentiation (Figure 4B, D,E). Further, multi-lineage differentiation potential was assessed and quantified using a two- dimensional directed differentiation assay. On average, all tested cell lines were capable to efficiently differentiate into cells of the mesodermal lineage (92.61% ± 5.99, mean ± SD), endodermal lineage (96.61% ± 2.30) and ectodermal lineage (88.35% ± 9.97), as determined by flow cytometry (Figure 4H-J). The results obtained from flow cytometry, including the expression of pluripotency-associated markers and the multi-lineage differentiation potential, were confirmed by representative immunofluorescence staining (Figure 4F, G). Based on phenotypic assessment, all iPSC lines met pluripotency-associated criteria (Table 3), showing no relevant intra-or inter-donor variability, as all cell lines performed equally well in aforementioned assays.

**Figure 4:**
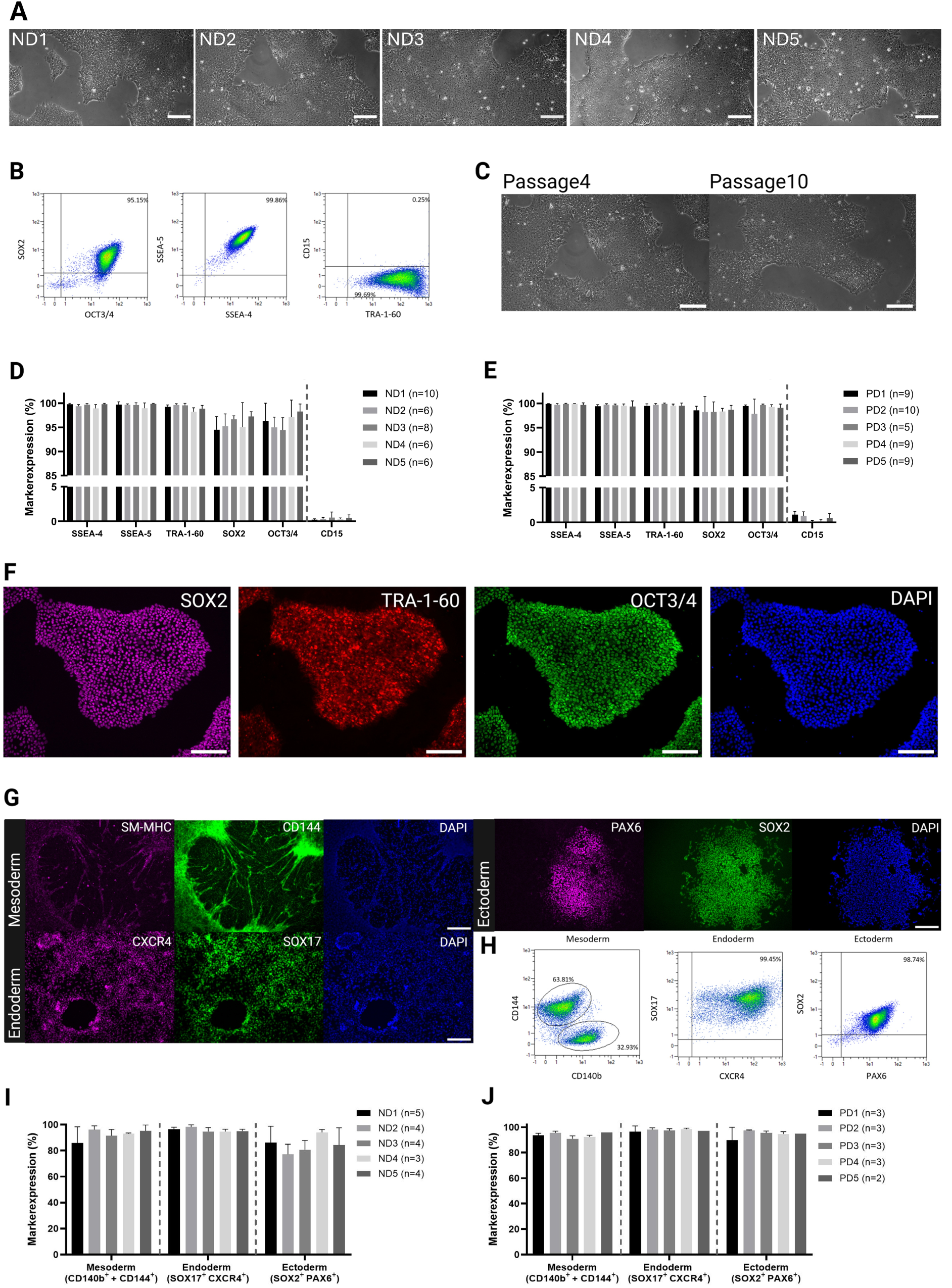
iPSC phenotype characterization. (A) Representative brightfield images of the morphology of derived iPSC lines in P4 from five non-diseased donors. (B) Gating strategy for the quantitative flow cytometry analysis of iPSCs for indicated pluripotency-associated markers and CD15 as an indicator of spontaneous differentiation. (C) Representative bright-field images illustrating maintenance of characteristic iPSC morphology over successive passages (passage four, ND2, same capture as in A). (D,E) Expression of pluripotency-associated markers (SSEA-4, SSEA-5, TRA-1-60, SOX2, OCT3/4) and one marker of spontaneous differentiation (CD15) of derived iPSC lines in P4 from five non-diseased donors (D) and five PD patients (E), as measured by quantitative flow cytometry analysis. Data are represented as mean + SD. (F) Representative immunofluorescence microscopy of iPSCs in P4 for pluripotency-associated markers (SOX2, TRA-1-60, OCT3/4). Nuclei were counterstained with DAPI. (G) Representative immunofluorescence microscopy of iPSCs in P5 differentiated into progenitors of the three germ layers mesoderm (SM-MHC, CD144), endoderm (CXCR4, SOX17) and ectoderm (PAX6, SOX2). Nuclei were counterstained with DAPI. (H) Gating strategy for the quantitative flow cytometry analysis of indicated markers after seven days of directed differentiation of iPSC into meso-, endo- and ectodermal cells. (I,J) Multi-differentiation potential of 3-5 iPSC lines per donor in P4 derived from five non-diseased donors (I) and five PD patients (J), as measured by quantitative flow cytometry analysis. Data are represented as mean + SD. Scale bars, 200 µm.

**Table 3:** Quality and acceptance criteria for iPSC line evaluation.

| Criteria | Method | Requirement to pass |
| --- | --- | --- |
| Morphology | Brightfield microscopy | PSC-like morphology:<br>high nuclear-to-cytoplasmic ratio,<br>prominent nucleoli, growth in tightly<br>packed colonies with well-defined<br>borders and absence of spontaneously<br>differentiated cells |
| Pluripotency-associated marker<br>expression | Quantitative flow cytometry | TRA-1-60: >85%<br>SSEA-4: >85%<br>SSEA-5: >85%<br>SOX2: >85%<br>OCT3/4: >85%<br>CD15: < 2.50% |
| Multi-lineage<br>differentiation<br>potential | Quantitative flow cytometry | Mesoderm (CD144+CD140b+): >60%<br>Endoderm (CXCR4+/SOX17+): >60%<br>Ectoderm (PAX6+/SOX2+): >60% |
| CNV analysis | SNP microarray | Absence of critical *de novo* CNVs |
| Cancer-associated<br>risk profile | Whole exome sequencing | Absence of pathogenic or likely<br>pathogenic *de novo* variants |

### Genetic integrity and cancer-associated gene risk profiling as key quality control filters excluded a subset of iPSC lines

There is consensus within the field that PSCs are susceptible to acquiring recurrent genetic abnormalities during prolonged culture (13,27,28), potentially rendering affected iPSC lines unsuitable for downstream applications. Therefore, genetic integrity needs to be checked during the initial qualification of iPSC lines and subsequently monitored at regular intervals across additional passages. In this study, genetic stability was first analyzed using the iCS-digital^TM^ Aneuploidy test, a digital 24-probe PCR that claims to detect more than 90% of recurrent numerical CNVs in PSCs (29). All iPSC lines derived from non-diseased donors (n=36) were analyzed. In 34 of 36 iPSC lines, no aberrations were detected at the examined loci, indicating absence of CNVs within these regions. One cell line revealed a loss in chromosome 5q, another cell line showed a gain in Xq, rendering both cell lines unsuitable for downstream applications (Figure S1A).

To achieve a more comprehensive genomic characterization, CNVs were screened using the next-generation Infinium Global Screening Array-24 v3.0 (Illumina), which offers broad genomic coverage and allows the detection of even small CNVs in the kilobase range. A call rate of >0.98 was achieved in all samples, thus passing the manufacturer’s recommended QC threshold (Figure S1B). For the iPSC lines from the non-diseased donor cohort, we were able to confirm the previously detected CNVs from the digital 24-probe PCR using the SNP microarray. Since the SNP array provides broader genomic coverage, additional aneuploidy screening by digital 24-probe PCR, which targets known PSC-associated hotspots only, was considered redundant and therefore not performed for the PD patient-derived iPSCs. We evaluated *de novo* CNVs detected relative to fibroblasts using the StemCNV-check pipeline (30). This pipeline facilitates all stages of analysis and includes specialized features such as sample-to-reference comparison, a CNV scoring system according to CNV biological impact and identity verification via SNP genotyping profiles, all tailored to human PSC. In total, 9/36 iPSC lines (25.0%) derived from the non-diseased donor cohort and 5/42 iPSC lines (11.9%) from the PD patient cohort were excluded from the study because of critical CNVs called by the StemCNV-check pipeline (Figure 5A,C). Notably, exclusion rates differed substantially between individual donors, ranging from 0% (PD3, PD4, and PD5) to 33.33% (ND4, ND5, PD1; Figure 5B,D). Identified CNVs were distributed across the genome without evidence of specific hotspots (Figure 5E,F). Sample-to-reference comparison allowed the exclusion of inter-donor contaminations, as each iPSC line could be unambiguously matched to its parental donor fibroblast line, thereby confirming their genetically correct ancestry.

**Figure 5:**
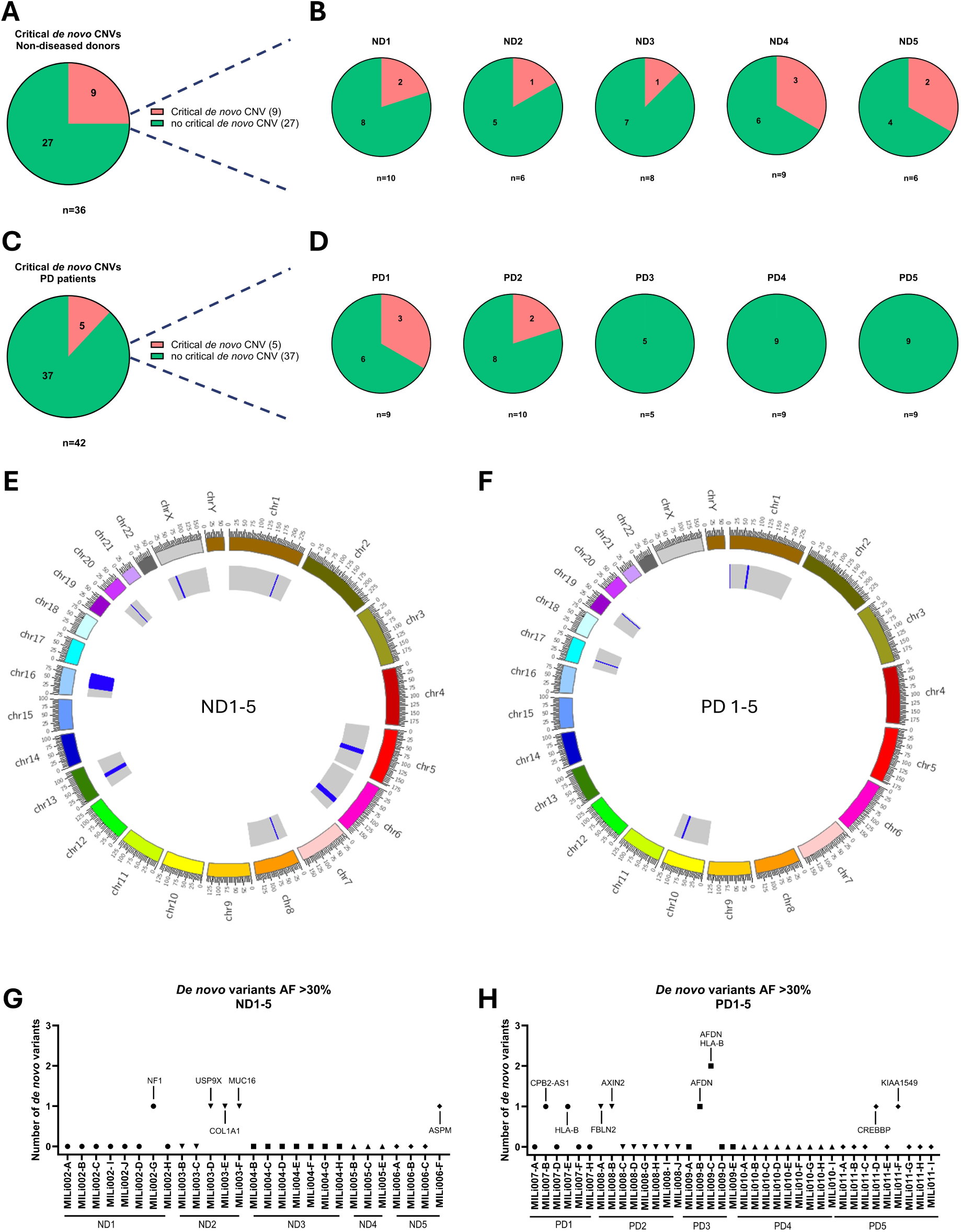
Genetic integrity and cancer-associated risk profile. (A,B) Pie charts illustrate the proportion of cell lines with (red) and without (green) critical *de novo* CNVs, displayed cumulatively across all non-diseased donor-derived iPSCs (A) and stratified by individual donor (B). (C,D) Pie charts illustrate the proportion of cell lines with (red) and without (green) critical *de novo* CNVs, displayed cumulatively across all PD patient-derived iPSCs (C) and stratified by individual PD patient (D). (E,F) Circos plots illustrating the genome-wide distribution of critical *de novo* copy number variations (CNVs) identified in iPSC lines derived from non-diseased donors (E) and PD patients (F). Within the inner circle, the entire chromosome segment is highlighted in grey, and the blue bars indicate the position of the critical CNV on the chromosome. (G,H) Number of QCI-predicted *de-novo* pathogenic or likely pathogenic cancer-associated variants according to ACMG/AMP guidelines. Only pathogenic and likely pathogenic variants with an AF >30% (fixed variants) from non-diseased donors (G) and PD patients (H) were considered. AF: allele frequency

For oncogenic risk assessment, we selected the subset of iPSC lines that had successfully passed prior QC assays for whole-exome sequencing (WES). The analysis focused on the detection of pathogenic and likely pathogenic variants in cancer- associated genes, employing the Clinical Insight (QCI) Interpret platform (QIAGEN) to ensure comprehensive and biological interpretation. We investigated variants in 693 cancer-associated genes and PSC hotspots. The 200X sequencing coverage allowed to reliably call variants with an allele frequency (AF) of >5% (Table S3,4). Given our primary focus on generating genetically unaltered iPSC lines without increased oncogenic potential, our analysis was restricted to *de novo* variants in cancer-associated genes (OnkoKB, COSMIC) as well as genes specifically selected for their relevance to stem cell biology (Stem-Seq™ (31)). Following the criteria established by Popp et al. 2018 (32), we considered fixed variants with an AF >30% as an exclusion criterion. Notably, most variants that could be detected were already present in the fibroblast parental cell line. However, 5/27 of the non-diseased donor cohort (Figure 5G) and 8/37 of the PD patient cohort (Figure 5H) acquired pathogenic or likely pathogenic *de novo* variants rendering them unsuitable for further use.

### Bona-fide iPSC lines could be nominated for each donor

Having characterized the pluripotency-associated phenotype, the genetic integrity and the oncogenic risk profile of our candidate iPSC lines from both cohorts, bona-fide iPSC lines could be nominated (Figure 6A,B) for each donor. No cell lines were excluded due to an inappropriate pluripotency-associated phenotype, as all cell lines exhibited the desired morphology (78/78), marker expression (78/78), and multilineage differentiation potential (34/34). Notably, not all lines were assessed for multi-lineage differentiation potential due to the considerable workload, however the acquired data provide strong evidence that the untested iPSC lines are likely to fulfill this requirement. Genetic integrity assessment using SNP array-based CNV analysis, alongside oncogenic risk profiling through WES, led to the exclusion of a subset of iPSC lines. Here, we observed inter-donor differences with varying numbers of cell lines being excluded per donor. In the gradual selection process, only iPSC lines that passed each preceding QC assay were retained for further analysis. Ultimately, of the 78 generated iPSC lines, 51 (65.38%) met all predefined QC criteria and were classified as bona-fide. Importantly, at least one bona-fide iPSC line for each donor could be identified as a candidate for further downstream applications. Considering the cumulative dataset across all iPSC lines and integrating every QC layer, including the passing score for the (i) PSC phenotype assessment (78/78, 100%), (ii) SNP array-based CNV analysis (64/78, 82%), and (iii) cancer-associated risk profiling (51/64, 80%), we calculated that on average screening three cell lines provides a ≥95% probability of identifying at least one cell line that fulfills all QC criteria. This result is based on the following calculation, which considers the passing score of each consecutive QC layer: (1 − (1 − 0.82 × 0.80)³) = 0.96. Due to observed inter-donor variability, this calculation was applied individually for each donor, resulting in a required number of initial iPSC lines ranging from one (PD4) to eight iPSC lines (ND2) to ensure a ≥95% probability of identifying at least one iPSC line that fulfills all QC criteria (Figure 6C). For future sample size planning, we therefore conservatively estimated a lower bound of eight iPSC lines per donor as the minimum requirement to achieve at least one bona-fide iPSC line for downstream applications.

**Figure 6:**
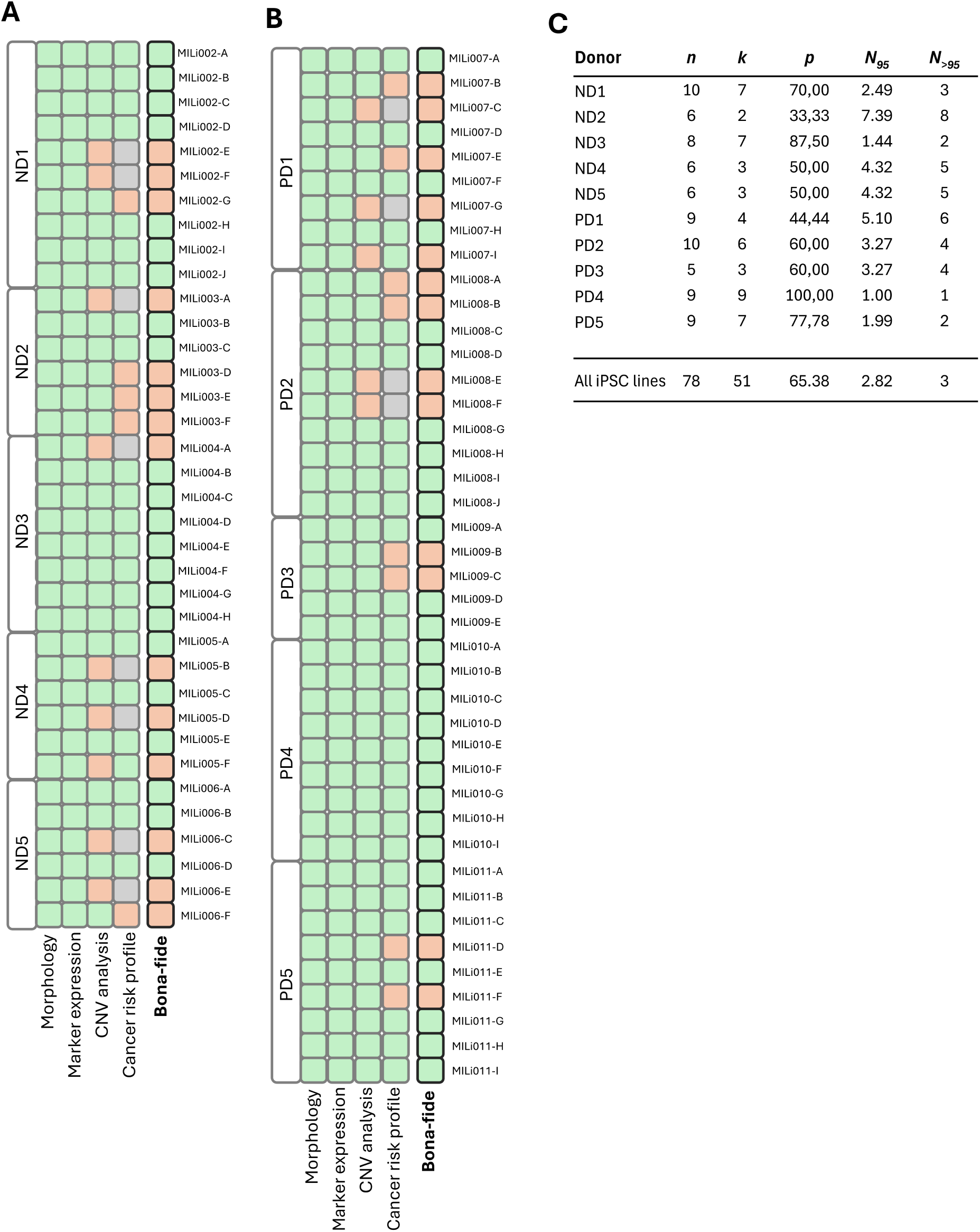
Individual iPSC line evaluation and selection of bona-fide iPSC lines. (A,B) Test results for QC criteria morphology, marker expression, CNV analysis, and cancer risk profile for non-diseased donors (A) and PD patients (B). Each iPSC line per donor was tested and evaluated individually. Green: passed; red: failed; gray: not tested (previously failed). (C) Success rates (*p*) were calculated for each experimental group (ND1-ND5, PD1-PD5) as the proportion of iPSC lines passing all sequential test criteria: *p=k/n* where *k* denotes the number of successfully passing iPSC lines and *n* the total number of tested iPSC lines for each individual donor. The number of iPSC lines required to obtain at least one successful clone with a target probability of 95% (*N_S5%_)* or greater than 95% and (*N_>S5%_)* was calculated using a Bernoulli model, as described in the Methods section.

### Automated and closed systems can replace manual and error-prone work in the iPSC manufacturing process

iPSC culture is still very much characterised by manual work governed by subjective decision that require skilful and trained personnel (e.g. for clonal picking of iPSC colonies). From a manufacturing perspective, automating subprocesses (Figure 7A) has the potential to streamline workflows and improve procedure consistency. mRNA-based reprogramming requires repetitive manual operator actions for transfections, media changes and harvesting. However, these processes can be automated on the CliniMACS Prodigy^®^. Proof-of-concept for successful reprogramming was demonstrated in two independent experimental runs using two different fibroblast cell lines (Figure 7B). Limitations inherent to manual iPSC colony picking, including labor-intensive handling and operator-dependent selection bias, can be effectively overcome through the application of microchip-based single-cell sorting of reprogrammed cultures using the MACSQuant^®^ Tyto^®^. This method preserves high cell viability after sorting (Figure 7C) and enriches TRA-1-60+ cells to >85%, independent of their initial target frequency (Figure 7D). Automated and gentle single-cell seeding was achieved with the VIPS™ Single Cell Seeding System, which provides image documentation to verify monoclonality. An established iPSC line, generated using the procedure described above, was selected as a representative example and used to seed five 96-well plates. An average of 73 ± 3.67 single cells (mean ± SD) were seeded per 96-well plate, of which approximately half formed colonies (Figure 7E). This corresponded to a colony-forming efficiency ranging from 46% to 53% across plates (Figure 7F), with an overall mean efficiency of 50.23% ± 3.16 (mean ± SD), demonstrating high consistency among all five plates. Compared to manual limiting dilution, VIPS^TM^-based seeding generated a larger pool of iPSC candidates for monoclonal outgrowth. For further iPSC expansion of cryopreserved or fresh iPSC, the CliniMACS Prodigy^®^ enables GMP-compliant, automated iPSC expansion in a closed and controlled system. iPSCs (MILi001-A) expanded using the CliniMACS Prodigy^®^ exhibited a proliferation capacity comparable to that of manually expanded iPSCs (Figure 7G), while maintaining high expression of pluripotency-associated markers (Figure 7H).

**Figure 7:**
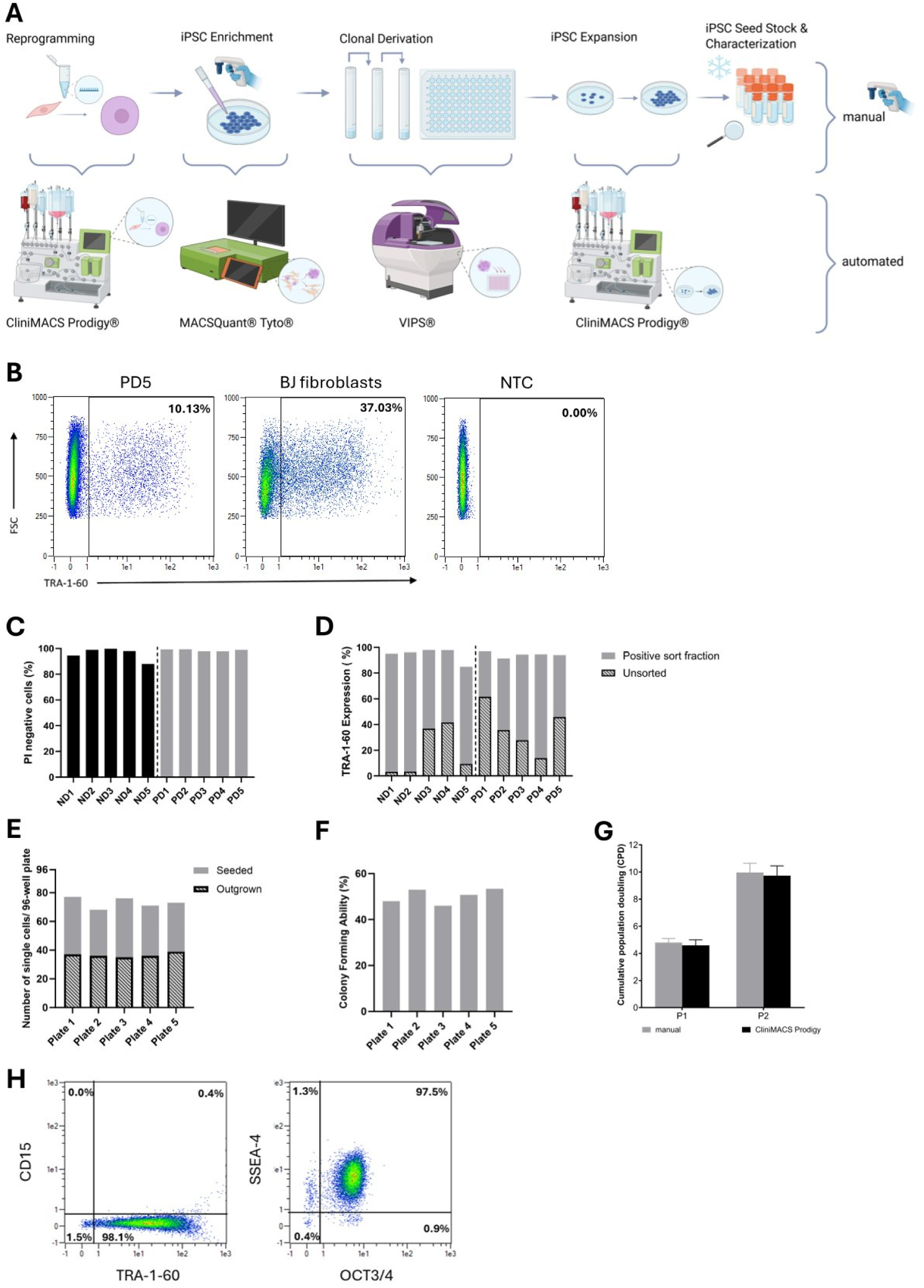
Transfer to automated platforms. (A) Schematic representation of the transfer from manual work to partially semi-automated and/or closed processes. (B) TRA-1-60 expression (%) after reprogramming PD5 fibroblasts (left) and BJ fibroblasts (middle) using the CliniMACS Prodigy® from two independent experimental runs. NTC (right): Non-transfected control. (C) Viability of cells from the positive sort fraction for five non-diseased donors (left) and five PD patients (right). Viability was assessed by PI staining; bars represent the percentage of viable (PI-negative) cells. (D) Sort performance showing the initial TRA-1-60-expression after reprogramming (dashed) and the Tra-1-60 expression of the positive sort fraction after microchip-based sorting (solid). (E) Number of wells with a single cell deposited in a 96-well plate by the VIPS^®^ (solid), and the Number of wells seeded with a single cell that showed an iPSC colony on day seven after seeding (dashed). (F) Colony-forming ability (%) of iPSC over five subsequently seeded 96-well plates (E) Colony-forming ability was defined as the capacity of single cells to proliferate and form colonies *in vitro*. (G) Cumulative population doublings (CPDs) of passages one and two of CliniMACS Prodigy®- expanded and manually-expanded iPSCs (MILi001-A) from three independent experiments. Data are presented as mean + SD. (H) Representative expression of pluripotency-associated markers (SSEA-4, TRA-1-60, OCT3/4) and CD15 as an indicator of spontaneous differentiation of CliniMACS Prodigy®-expanded iPSCs as measured by quantitative flow cytometry analysis.

## Discussion

We established and optimized a comprehensive workflow for the generation and selection of iPSC lines for autologous cell therapies. This end-to-end workflow entails fibroblast isolation straight from skin, reprogramming, iPSC enrichment, monoclonal derivation and QC to select bona-fide iPSC lines for downstream applications. The workflow was established using skin biopsies from five non-diseased donors aged 50-62 years and subsequently applied to relevant patient material from five PD patients aged 49-81 years, resulting in 78 monoclonal iPSC lines from ten donors.

Despite aging being a known limiting factor for reprogramming efficiency (33,34), we successfully reprogrammed iPSCs from all donors regardless of age and sex, however with varying degrees of efficiency. PD patients (male only) exhibited higher reprogramming efficiencies than non-diseased donors (predominantly female), which we attribute primarily to variations in sample processing rather than donor status. Notably, samples from PD patients were processed immediately (<2h post-biopsy), whereas samples from non-diseased donors were in transit for at least 48h due to logistical constraints. This prolonged transit time likely compromised cell quality and reprogramming efficiency. For the donors for whom reprogramming efficiency was determined (ND5, PD1-PD5), the mRNA-based reprogramming method reached reprogramming efficiencies exceeding 6.58%, outperforming other non-integrating methods such as Sendai virus and protein reprogramming (Sendai virus, 0.01-1% (6); protein, 0.001% (6)).

Exogenous mRNA degraded progressively, with no detectable residuals by passage one, while endogenous pluripotency gene expression was high, indicating a self-sustaining pluripotent state independent of exogenous factors. Thereby, generated iPSC lines were ready for banking, molecular characterization and further differentiation at a very early stage, minimizing culture duration and thereby reducing the risk of cultivation-induced aberrations (11,35). In contrast, non-integrative methods such as Sendai virus- or episomal-based reprogramming show a slower, passage-dependent decline of exogenous residuals. For instance, Schlaeger et al. (6) reported SeV RNA persistence in 100% of iPSC lines at P1-5, 53.8% at P6-8, and 21.2% at P9-11, while EBNA1 DNA was detected in 39.1%, 37.9%, and 33.3% of episomal iPSC lines at the same passage ranges. The resulting need for extended screening prolongs time in culture, increasing costs, workload, and the risk of aberrations, thereby complicating GMP implementation.

All iPSC lines consistently expressed pluripotency-associated markers at high levels and tested iPSC lines could successfully differentiate into cell types of the three germ layers, thus fulfilling ISSCR guideline standards (23,36). We demonstrated low inter- and intra-donor variability in terms of their pluripotency-associated phenotype, which implies that by keeping factors like somatic cell type, reprogramming method, and culture platform constant, standardization of the process and comparability of the cells is highly favoured, avoiding high donor-to-donor inconsistencies.

PSC populations are prone to acquire recurrent CNVs during reprogramming and extended cultivation (37–40). To assess the CNV burden, we first performed iCS-digital^TM^ Aneuploidy PCR analysis targeting recurrent PSC hotspot regions. Among 36 iPSC lines from non-diseased donors, only two exhibited CNVs in recurrent hotspot regions, indicating that early passages preserve genomic integrity before aberrant clones can expand. For deeper analysis we performed genome-wide SNP microarray profiling with subsequent CNV evaluation using the StemCNV-check pipeline (30). Conventional CNV analyses often rely on size thresholds (32,41), neglecting effects on coding regions or application-specific requirements. Depending on the chosen size threshold, varying numbers of cell lines may be excluded, raising the general question which threshold is appropriate for CNV annotation in PSCs. Rather than excluding iPSC lines only based on CNV size, we performed individualized assessments of CNVs. The StemCNV-check pipeline applied here focuses on genomic regions involved in cell cycle regulation, apoptosis, and proliferation - key pathways prone to selective advantage in PSCs. Given the wide spectrum of diseases for which autologous cell therapies are considered, the pipeline can be tailored to specific disease-related gene regions in the future. It is noteworthy that two out of five PD patients (PD2, PD5) carried clinically relevant *GBA* variants. Although this does not affect the usability of these specimens for our study, these patients would not benefit from autologous therapy without additional gene correction. Alternatively, they may qualify for a potential allogeneic treatment regimen. Of the 78 iPSC lines analyzed, 14 exhibited critical CNVs that were either absent or present only at undetectable levels in the parental fibroblasts. Cumulatively, less of the tested PD iPSC lines (5/42, 11.90%) than non-diseased donors (9/36, 25.00%) showed critical CNVs, maybe as the upper thigh (PD) allowed sampling from a relatively sun-protected biopsy site compared to the abdomen (ND). Most CNVs were small (∼500 kb), falling below the lower detection limit of 5-10 Mb typically assumed for G-banded karyotyping. Consequently, many CNVs identified here, would escape detection by conventional methods, risking the use of genomically unstable iPSC lines in clinical applications when relying solely on superficial profiling techniques like G-banding. However, although SNP microarrays enable the detection of the most common genomic alterations in iPSCs, they are unable to identify balanced chromosomal rearrangements. Therefore, G-banding could be additionally added to confirm structural integrity for final candidate iPSC lines. It is important to note that objective genetic QC criteria are still lacking, primarily due to the complexity of interpreting whole-genome sequencing data in the context of risk profiling.

A fundamental criterion for future clinical grade iPSC lines also includes absence of additionally acquired potentially harmful mutations (e.g. reported cancer-related mutations). In the context of autologous iPSC therapy, the aim is therefore to generate iPSC lines derived with minimal genetic divergence from the donor’s genetic background (32), why we focused on cancer-associated and PSC-relevant *de novo* variants as an exclusion criterion for downstream use. We could confirm literature findings that mutational burden mostly derives from starting material (42,43). While iPSC lines largely retained the genetic background of their parental cells, 16 sporadic critical *de novo* CNVs and 14 oncogenic variants arose, likely due to mutagenesis or selection during reprogramming or monoclonal derivation underscoring the necessity of individual assessment before downstream use. Our data suggest that applying phenotypic selection criteria does not lead to the exclusion of cell lines from the candidate pool. In contrast, genetic integrity and cancer-associated risk profiles were more selective.

We propose the following QC pipeline for selecting bona-fide iPSC lines: (i) Morphological assessment and flow cytometry-based analysis of pluripotency-associated markers. (ii) SNP array-based genetic integrity analysis, owing to its cost-effectiveness and high resolution. (iii) WES for oncogenic risk profiling. (iv) Multi-lineage differentiation assay. Although the multi-lineage differentiation assay, has proven to be non-selective in our mRNA-based approach, it remains a necessary step to formally demonstrate pluripotency (36). Thus, our pipeline addresses the core aspects of ISO 24603:2022 and ISSCR standards 2023 for fundamental iPSC characterization, including cell line authentication, assessment of pluripotency, and evaluation of genetic integrity using recommended assays. Nevertheless, further clarification regarding specific methodological parameters (sequencing depth, CNV size thresholds) and the alignment in the stringency of selection criteria remains to be discussed across laboratories in the field.

Based on our dataset, generating three iPSC lines yielded a >95% probability of obtaining at least one bona-fide iPSC line (cumulative dataset). For comparison, Popp et al. 2018 conducted a comprehensive study including >70 iPSC lines as part of the ForIPS consortium (32) and concluded that on average eight iPSC lines (cumulative dataset) are required to have a ≥90% chance to get at least one cell line that passed all QC criteria, when considering conventional karyotyping, CNV analysis and oncogenic risk profiling by WES. However, it should be noted that the authors applied a threshold of ≥100 kb for CNV annotation without individual biological judgement and analytical methods for WES differed. We noticed in our study that exclusion rates varied across donors. Thus, in the best-case scenario, only one cell line was needed to achieve a >95% probability of obtaining at least one bona-fide iPSC line, while in the worst-case scenario, eight were required. These donor-specific differences likely reflect variations in fibroblast quality, genetic predisposition, or sporadic events originating from the starting material. Although the cumulative number of iPSC lines analyzed was substantial (n= 78), the number of iPSC lines per donor remained limited (n= 5-10), introducing a degree of statistical uncertainty into the estimated success probabilities. Nevertheless, these estimates provide a useful framework for approximating the number of iPSC lines required in comparable experimental settings and may support study design and resource planning. Accordingly, the calculated sample sizes should be taken as practical guidelines rather than strict requirements. Future studies incorporating larger datasets across multiple experiments could further improve the robustness of success probability estimates and enable the application of more advanced statistical modeling approaches.

Having generated multiple iPSC lines, we can summarize that the total duration for the generation of monoclonal iPSC lines spans approximately seven weeks. The process can be delineated as follows: (i) fibroblast isolation, one week; (ii) reprogramming, two weeks; (iii) iPSC expansion, four weeks. As QC analyses are performed on cryopreserved cell stocks post-thawing, they do not contribute to the cumulative time cells are kept in culture. Monoclonal iPSC expansion is the most time-consuming step in the workflow; however, this additional time requirement is accepted to enable highly selective screening and to minimize uncertainties in e.g. genetic integrity associated with polyclonal iPSC lines.

Toward manufacturing and GMP-compliant transfer, we provided proof-of-concept to transfer processes like reprogramming and monoclonal derivation to semi-automated and/or closed systems to enhance standardization and product safety through reduced contamination risk. Recently, Shimizu et al. 2026 demonstrated successful Sendai-based reprogramming of PBMCs using the CliniMACS Prodigy^®^ (44). In this study, we expanded the applicability of the CliniMACS Prodigy^®^ to mRNA-based fibroblast reprogramming. Nevertheless, the establishment of a fully integrated end-to-end, automated, and closed manufacturing process remains under development.

In summary, we present a robust and efficient workflow enabling generation of human monoclonal iPSCs from skin punch biopsies from each donor within seven weeks. A systematic QC cascade ensures selection of high-quality iPSC lines suitable for downstream applications. Thus, this study contributes to the advancement of personalized iPSC-based research and future translational efforts.

## Resource availability

### Lead contact

Requests for further information or resources associated with this study should be directed to Sebastian Knöbel.

### Materials availability

This study did not generate any new or unique reagents.

### Data and code availability

Flow cytometry data have been deposited in FlowRepository. Whole exome sequencing and SNP array data have been submitted to the European Genome-phenome Archive (EGA): Accession numbers will be publicly available as of the date of publication. Any additional information required to reanalyze the data reported in this paper is available from the lead contact upon request.

## Acknowledgements

This work was supported by funding from BMBF (iPStemRNA, 01EK1610C). The authors would like to thank Sebastian Knöbel’s entire group for their support and collaboration.

## Author contributions

Conceptualization: SK, AB, HJ. Methodology: DH, CW, HJ, UW, CR, TK, SB, CL. Investigation: DH, CL. Formal Analysis: DH, CW. Visualization: DH. Supervision: SK, HJ. Writing - Original Draft: DH. Writing - Review & Editing: All authors.

## Methods

### Ethical concerns

Written informed patient consent was obtained from all PD patients with approval of the Ethical committee of the University Bonn (Ref. No. 311/14) for the collection of a skin punch biopsy. Skin punch biopsies from non-diseased controls were obtained from surgery leftovers mediated via a commercial provider (Biopredic International, Azenta Life Sciences). Tissue collection and distribution were conducted under approval from the French Ministry of Research (N° AC-2023-5431). Volunteers gave written consent before. The samples were received without patient identifiers, only providing information about age, sex and ethnicity. Biological material was handled following required safety procedures. All donors were serologically screened negative for HIV-1, HIV-2, HBV, HCV prior to sample processing. All donors were self-reported Caucasian. Ethnicity was not considered a study variable and is not anticipated to affect the outcomes presented here.

### Tissue processing of skin punch biopsies

Fibroblast-derived iPSCs were generated from non-diseased donors using abdominal skin leftovers from plastic surgeries. For the generation of fibroblast-derived iPSCs from PD patients a 4 mm^2^ skin punch biopsy was taken from the upper part of the thigh. For transportation between sites, skin tissue was kept in MACS^®^ Tissue Storge Solution (130- 100-008, Miltenyi Biotec). Given the increased susceptibility of skin-derived samples to microbial contamination, sample shipment, cell isolation and initial fibroblast expansion were conducted in the presence of anti-bacterial and anti-fungal reagents (A5955, Sigma Aldrich). However, these were removed from the medium prior to reprogramming. The mechanical and enzymatic dissociation of skin tissue was performed using the human Whole Skin Dissociation Kit (130-101-540, Miltenyi Biotec) according to manufacturer’s instructions. Isolated cells were stained with CD90-VioBlue^®^ (1:50) in PEB (PBS, 2 mM EDTA, 0.5% BSA) for 10 min at 4 °C and the CD90+ frequency was measured using the MACSQuant® flow cytometer. After tissue dissociation, 10,000 CD90+ cells/cm² were seeded onto LN-coated (0.5 µg/cm^2^; LN521-05, BioLamina) in supplemented MSC-Brew GMP medium (170-076-332, Miltenyi Biotec) with regular media changes every 3-4 days. Fibroblasts were maintained in culture for around one week (until confluency was reached) to start reprogramming and to generate a low-passage cell stock for cryopreservation. No matrix was used for further fibroblast expansion. Before reprogramming, fibroblasts were tested for mycoplasma using the MycoStrip® 2.0 - Mycoplasma Detection Kit (rep-mysv2-10).

### mRNA-based reprogramming of primary fibroblasts

Dermal fibroblasts were tested negatively for mycoplasma before reprogramming. On day-3 fibroblasts were seeded on Laminin521-coated plates (0.5 µg/cm^2^) with a density of 1,500 cells/cm^2^-3,000 cells/cm^2^ in supplemented MSC-Brew GMP medium at 37 °C, 5% CO_2_. The StemMACS^TM^ iPSC mRNA reprogramming Kit (130-132-990, Miltenyi Biotec) was used according to manufacturer’s instructions with minor modifications as specified in the following. Deviating from the provided protocol, fibroblasts were seeded and cultivated from day-3 to day0 in supplemented MSC-Brew GMP medium and StemMACS^TM^ PSC-Brew XF medium (130-127-865, Miltenyi Biotec) was used from day5 until the end of reprogramming. If fibroblasts were cryopreserved before reprogramming (PD1, PD4 C CliniMACS Prodigy^®^ reprogramming), fibroblasts were thawed in supplemented MSC-Brew GMP, expanded for five days and reprogrammed as described above.

For reprogramming using the CliniMACS Prodigy^®^ the Tubing Set 730 (200-073-623, Miltenyi Biotec) and the Adherent Cell Culture (ACC) software (version 1.1) were used. The chamber was coated with either 0.385 µg/cm^2^ MACSmatrix Laminin511 (130-136-454, Miltenyi Biotec, PD5) or with 0.55 µg/cm^2^ LN521 (BJ fibroblast). The mRNA-based reprogramming was carried out according to manufacturer’s instructions with minor modifications as specified in the following. The mRNA transfection complex was added to the StemMACS Repro-Brew medium, which was transferred to a bag and welded on valve 3 of the tubing set. From day5 on iPS-Brew GMP was used (170-076-317, Miltenyi Biotec). For the experiments presented here, 10% more cells were seeded compared to the small-scale setup.

### Microchip-based cell sorting for iPSC enrichment

Reprogrammed cells were harvested on day10 or day11 of the reprogramming protocol with TrypLE^TM^ (12563029, Thermo Fisher Scientific) and filtered using a 70 µm pre-separation filter (130-095-823, Miltenyi Biotec). Cells were stained with TRA-1-60-PE (Table S5, 1:50) for 10 min in PEB at 4 °C. Fluorescently labelled cells were resuspended in a suitable volume of MACSQuant^®^ Tyto^®^ Running Buffer (1.0 × 10^6^ cells/mL; 130-107- 206, Miltenyi Biotec) and loaded into the input chamber of a MACSQuant^®^ Tyto^®^ cartridge. The MACSQuant^®^ Tyto^®^ HS cartridge (130-121-549, Miltenyi Biotec) was used for non-diseased donors. The availability of a new Tyto^®^ cartridge developed for samples containing cells >20 µm diameter enabled the PD patient samples to be sorted in MACSQuant^®^ Tyto^®^ LS cartridges (130-133-260, Miltenyi Biotec), which turned out to be more suitable for reprogrammed cell cultures. After iPSC enrichment, sorted fractions were recovered from the cartridge under sterile conditions for downstream applications.

### Single cell seeding and iPSC cell line derivation

For manual single cell seeding of positive sorted cells, supplemented StemMACS^TM^ PSC- Brew XF medium was used in combination with the StemMACS^TM^ PSC-Support XF supplement (130-127-287, Miltenyi Biotec). To minimize the risk of depositing more than one cell per well, 0.3 cells/well were seeded in Laminin521-coated 96-well plates (1.5 µg/cm^2^). Media changes were performed as specified in the manufacturer’s protocol (StemMACS^TM^ PSC-Support XF). Cells were cultivated in 96-well plates for approximately 14 days, after which they were gradually expanded to larger culture formats: 96-well plate (P1), 24-well plate (P2), one well of a 6-well plate (P3), T75-flask (P4). Following the initial step in the 96-well format, cells were maintained according to standard iPSC culture practice (chapter iPSC culture). Once the cell lines reached 80-90% confluency in the 75- flask, they were cryopreserved to generate a low-passage iPSC line seed stock in P4. Quality control analyses were performed on the P4 seed stock. All generated iPSC lines are registered in the human Pluripotent Stem Cell Registry hPSC^reg^® (https://hpscreg.eu) (36) with corresponding registration IDs (Table 2).

For automated single cell seeding, the Solentim Verified in-situ plate seeding (VIPS^TM^) device was used according to manufacturer’s instructions (Advanced Instruments). iPSCs were diluted to 9,500 cells/mL in supplemented StemMACS^TM^ PSC-Brew XF medium in combination with the StemMACS^TM^ PSC-Support XF supplement and 0.5 µg/cm^2^ iMatrix511-E8 (AMS.892 011, amsbio). Following cell seeding, wells were filled with culture medium (StemMACS^TM^ PSC-Brew XF, StemMACS^TM^ PSC-Support XF supplement, iMatrix511-E8) to a final volume of 100 µL. Plates were imaged on the VIPS^TM^ two hours after seeding and on day one, seven, and 14 to confirm monoclonal outgrowth.

### Statistics

The success probability of generating bona-fide iPSC lines was estimated for each experimental group (ND1-ND5, PD1-PD5) as the proportion of iPSC lines that successfully passed all sequential QC criteria. The success rate *p* was calculated as:

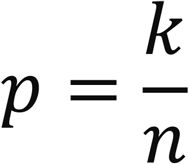

where *k* denotes the number of iPSC lines that passed all QC criteria and *n* the total number of tested iPSC lines per donor. The minimum number of iPSC lines required to obtain at least one bona-fide iPSC line with a target probability of 95% was calculated based on the Bernoulli model:

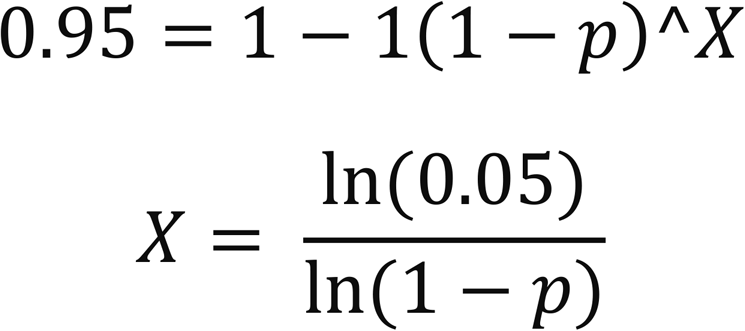

Given the limited number of observations per group, estimation uncertainty is substantial. Although confidence intervals could be used to derive conservative lower bounds, this would result in disproportionately large sample size requirements for downstream applications. We therefore report empirical success rates (*k*/*n*) and interpret them as approximate indicators rather than precise estimates.

### iPSC culture

iPSCs were maintained on Laminin521-coated plates (0.5 µg/cm^2^) in supplemented StemMACS^TM^ PSC-Brew XF medium in a humidified incubator at 37 °C, 5% CO_2_. For initial plating after thawing or passaging, the medium was supplemented with 2 μM StemMACS^TM^ Thiazovivin (130-106-542, Miltenyi Biotec) which was withdrawn from the cell culture after 48h. Thereafter, the medium was refreshed regularly every 24-48h. The cells were continuously monitored for bacterial and fungal contamination using a brightfield microscope. Upon reaching 80% confluency, cells were passaged using TrypLE^TM^ and replated as single cells at a density of ∼10,000 cells/cm^2^. iPSCs were cryopreserved in StemMACS^TM^ Cryo Brew with a density of 1.0 × 10^6^ cells/mL.

For iPSC cultivation using the CliniMACS Prodigy^®^ the Tubing Set 730 (200-073-623, Miltenyi Biotec) and the Adherent Cell Culture (ACC) software (version 1.4.30.184) were used. To compensate for process-related cell loss during automated cultivation, the culture vessels were coated with 10% more matrix (LN521; 0.55 µg/cm²) than in the corresponding manual small-scale cultures (0.5 µg/cm²). Cells were cultured in Cell Culture Units (CCUs) during passage 1 (P1) and transferred to Corning CellSTACK® culture chambers (3268, Corning) for passage 2 (P2). Manual small-scale cultures were performed in 6-well plates (P1) and T75-flasks (P2). iPSCs were maintained following routine culture procedures, including regular medium changes and passaging at appropriate confluency as specified above but using iPS-Brew GMP medium (170-076-339, Miltenyi Biotec) and 6,250 cells/cm^2^ cell density. Cell growth was evaluated over two consecutive passages.

### Assessment of iPSC differentiation potential

iPSCs were differentiated into cell types of the three germ layers using the StemMACS™ Trilineage Differentiation Kit (130-115-660, Miltenyi Biotec) according to the manufacturer’s protocol with minor modifications as specified below. Specifically, 1.5-fold the recommended cell number was seeded for each lineage. The multi-lineage differentiation potential was evaluated through quantitative flow cytometry and immunocytochemical analysis using lineage-specific antibodies (Table S5).

### Immunocytochemistry

Cells were fixed with 4% paraformaldehyde (J19943-K2, Thermo Fisher Scientific) for 10 min at room temperature prior to immunostaining. After blocking for 30 min in Blocking Buffer (10% FCS, 0.1% Triton X-100 in PBS), conjugated antibodies (Table S5) were diluted in Blocking Buffer and incubated overnight at 4 °C. Following a DPBS wash, 4′,6-diamidino-2-phenlindole (DAPI; 1 µg/mL; D1306, Thermo Fisher Scientific) was applied in Blocking Buffer and incubated for 45 min at room temperature. Fluorescence images were acquired using the EVOS^TM^ M5000 Imaging System with subsequent processing in ImageJ (45). Overview images of entire wells were obtained using the Cytation^TM^ 3 Cell Imaging Multi-Mode Reader. To determine reprogramming efficiency, overview images were blurred in ImageJ (Gaussian Blur) and the number of OCT3/4+ colonies was quantified using the segmentation tool integrated in the MACS^®^iQ View Analysis Software (version 1.3.1). Reprogramming efficiencies were calculated as follows: 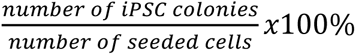

### Flow Cytometry

Flow cytometric data were acquired using the MACSQuant^®^ Analyzer 10 and the MACSQuant^®^ Analyzer 16 in combination with the MACSQuantify^TM^ Analysis Software (version 2.13.0). Cells were distinguished from debris based on forward scatter (FSC) and side scatter (SSC) properties. Dead cells were excluded after addition of Propidium Iodide (130-093-233, Miltenyi Biotec) to the single cell suspension (1 µg/mL). Surface antibody staining was performed in PEB Buffer for 10 min at 4 °C with fluorescent-conjugated antibodies using recommended concentrations (Table S5). Following incubation, cells were washed with PEB and centrifuged at 300 x g for 5 min. After surface marker staining, intracellular epitopes were stained using either the FoxP3 Buffer Set (130-093-142, Miltenyi Biotec) or the Inside Stain Kit (130-090-660, Miltenyi Biotec) according to provided protocols. To adjust laser voltages and compensate for spectral overlap, single antibody-stained samples and unstained samples were used.

### Quantitative Real-time PCR (RT-qPCR)

Total RNA was extracted using the RNAeasy Mini Kit (74104, QIAGEN). RT-qPCR was performed using the Takyon^®^ SYBR MasterMix (#UF-NSMT-B0701, Eurogentec) in combination with the Takyon^®^ One-Step Kit Converter (#UF-RTAD-D0701, Eurogentec) on 100 ng total RNA per reaction, in the 7900 HT Fast Real-Time Thermo Cycler. Amplification was carried out in 20 µL total reaction volume with 40 cycles of a 95 °C denaturation phase, followed by a 63 °C annealing phase and a 72 °C elongation phase. For detection of endogenous and exogenous RNA during reprogramming the probe-based detection Mastermix Takyon^®^ No Rox ProbeMaster Mix dTTP blue (UF-NPMT-B0701, Eurogentec) was used in combination with the Takyon^®^ One-Step Kit Converter and the designed probes. Exogenous and endogenous transcripts were distinguished based on sequence differences in their coding and untranslated regions. Primer and probe concentrations were titrated for each gene (Tab. S4). Gene expression analysis was performed using the 2^-ΔCT^ method (46) by normalizing relative gene expression to the transcription levels of the housekeeping gene.

### iCS-digital^TM^ Aneuploidy PCR

Cryopreserved cell pellets (1 × 10⁶ cells) were submitted to Stem Genomics (France) for iCS-digital^TM^ Aneuploidy PCR testing to detect recurrent genetic abnormalities in iPSCs (29). The supplier provided evaluated reports.

### SNP array-based CNV Analysis

DNA was extracted using the DNeasy Blood & Tissue Kit (#69504, QIAGEN). Genotyping was performed at the LifeCBrain Genomics Service Center (Bonn, Germany) utilizing the Infinium™ Global Screening Array-24 + Multi Disease Content v3.0 BeadChip (Illumina). A Copy Number Variation (CNV) analysis was performed using the StemCNV-check pipeline (version 0.7.3). The StemCNV-check pipeline (30) utilizes the genome-wide distributed SNP data of the 730.059 specifically designed markers, to run two independent algorithms for CNV detection: Circular Binary Segmentation (CBS) and PennCNV. The resulting CNVs, specifically *de novo* CNVs compared to the reference sample, are then ranked according to the Check-Score: A measure of the copy number (CN) and length of the CNV as well as its overlap with published stem cell hotspot, dosage sensitivity, and cancer-genes. Putative CNV calls (either marked as “critical”, due to stem cell hotspot overlap, or as “reportable”, due to overlap with dosage sensitivity or cancer genes) were manually reviewed to confirm validity (Table S1, S2).

### Whole Exome Sequencing (WES)

Genomic variants were identified via CLC Genomics Workbench Software (version 23). The variant identification template workflow “Identify QIAseq Exome Germline Variants” was adapted covering, among other processing steps, the functions ”Structural Variant Caller” and ”Low Frequency Variant Detection”. Resulting SNPs and indels were exported and further imported into QCI Interpret software (QCII). QCII was used for further filtering (Filters: “CallQual>20; PassedUpstream; AF>5%; CommonVariants; Predicted Deleterious”) and annotation of variants (Diagnosis: “Cancer”). Additional downstream processing using R (version 4.4) was applied to combine samples from multiple QCII analyses and to consider further custom filtering. Only variants in genes known to be relevant (COSMIC (48), OnkoKB (48,49), Stem-Seq™(31)) and annotated as “pathogenic” or “likely pathogenic” were kept. Variants were classified in accordance with guidelines of the American College of Medical Genetics and Genomics and Association for Molecular Pathology (ACMG/AMP) (50). The following additional criteria were assessed and subsequently applied to exclude variants: (i) Variants shared with fibroblasts samples, (ii) Variants with relative frequency <30% in all samples and no variant previously reported (i.e., present in ClinVar or COSMIC), (iii) Variants in the genes *TDG*, *TYRO3* and *MUC4* due to presence of many unspecific variants in all samples. The final set of variants was further manually assessed based on information gathered during all processing steps including read mapping and annotation of known variants.

### MACSima imaging cyclic staining (MICS)

BJ fibroblasts (CRL2522™, ATCC^®^) were reprogrammed as described above on Laminin521-coated (1.0 µg/cm^2^) MACSwell^TM^ Imaging plates (130-124-677, Miltenyi Biotec). On day11 cells were fixed and pre-stained with DAPI (1 µg/mL) following the MACSima™ sample preparation protocol for adherent cells using the MACSima^TM^ Stain Support Kit (130-127-574, Miltenyi Biotec). iPSC colonies and regions of interest (ROIs) were selected based on the DAPI signal specifically targeting areas with densely packed, round-shaped nuclei. The MACSima™ system operates through iterative fluorescence staining, image acquisition, and signal erasure, using multiple fluorochrome-conjugated antibodies per cycle (51). Conjugated antibodies (Table S5) were diluted in MACSima^TM^ Running Buffer and sorted in Deep Well Plates according to their epitope binding site. Surface antigens were stained first. Prior to intracellular staining, cells were permeabilized with Inside Perm in one cycle for 10 min. Images were generated using the MACSima^TM^ instrument with 10 min cycle length and analyzed using the MACS^®^iQ View Analysis Software (version 1.3.1, Miltenyi Biotec).

## Data availability, analysis and statistics

Flow cytometry data have been deposited in FlowRepository. Whole exome sequencing and SNP array data have been submitted to the European Genome-phenome Archive (EGA). Accession numbers will be provided upon assignment and included prior to publication. Raw data were analyzed and plotted using Microsoft Excel, GraphPad Prism 9, and BioRender. Genome-wide CNV distribution (circos plot) was visualized using SRplot (https://www.bioinformatics.com.cn/en), a freely accessible online platform for scientific plotting (52). Statistical tests used (GraphPad Prism 9) are indicated within respective figure legends.

## Supplemental

Table S1 StemCNV-check call evaluation ND1-ND5

Table S2 StemCNV-check call evaluation PD1-PD5

Table S3 Whole exome sequencing ND1-ND5

Table S4 Whole exome sequencing PD1-PD5

**Supplementary Figure 1:**
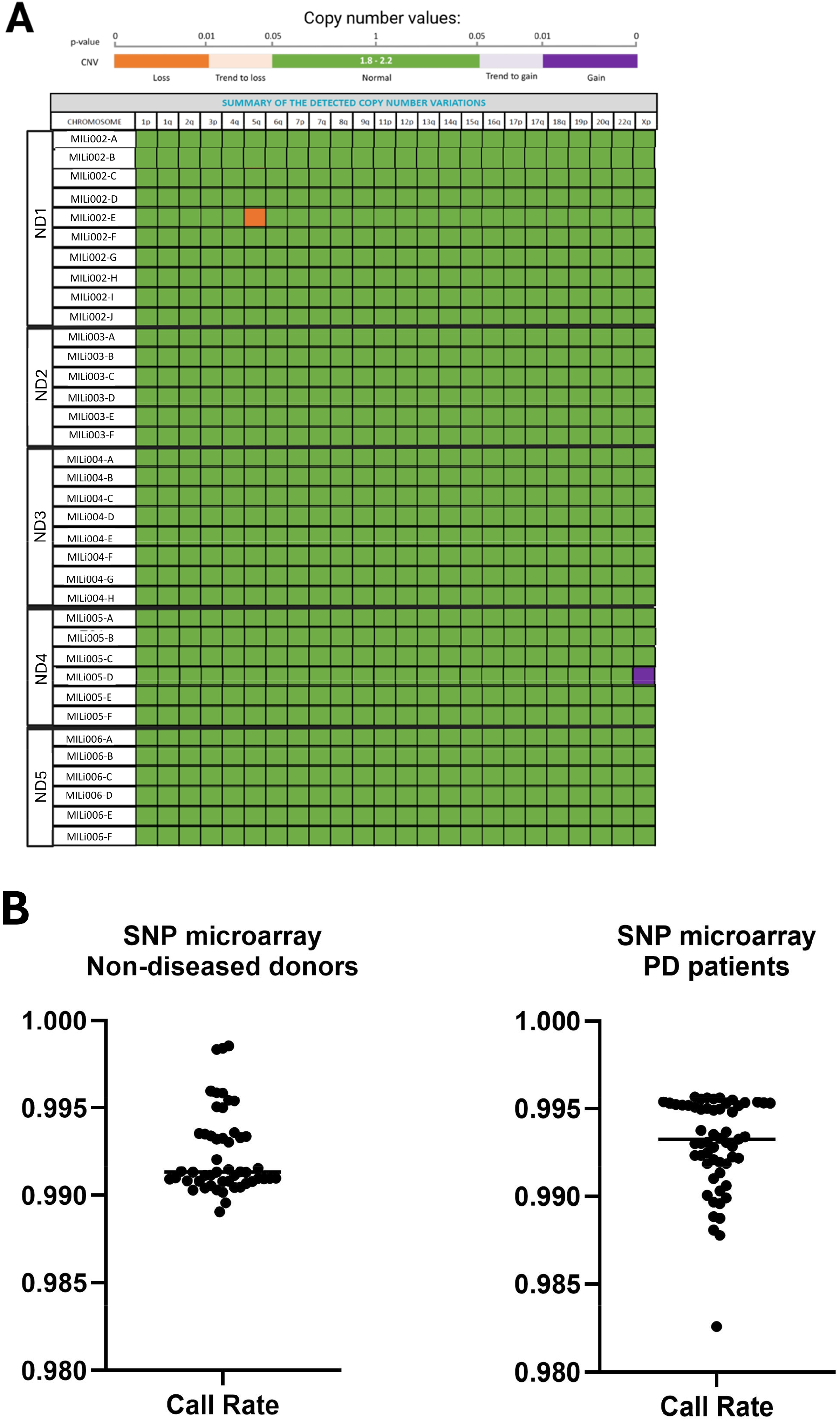
Genetic integrity. (A) iCS-digital^TM^ Aneuploidy PCR analysis of the most recurrent hotspots of CNVs for PSCs on each chromosome. Green corresponds to a normal diploid chromosome set (1.8-2.2). CNVs are categorized as trend to gain/loss (light purple/orange; p-value 0.01 < p < 0.05) or gain/loss (purple/orange; p-value < 0.01). (B) The call rate indicates the proportion of SNPs successfully genotyped per sample for iPSCs derived from non-diseased donors (left) and iPSCs derived from PD patients (right). The threshold for acceptable sample quality is ≥0.98.

**Table S5:** List of antibodies.

| **Antibody** | **Clone** | **Dilution** | **Order number** | **Vendor** |
| --- | --- | --- | --- | --- |
| **Flow cytometry** |  |  |  |  |
| CD90-VioBlue® | REA897 | 1:50 | 130-114-866 | Miltenyi Biotec |
| TRA-1-60-PE | REA157 | 1:50 | 130-122-921 | Miltenyi Biotec |
| CD15-PE-Vio®770 | VIMC6 | 1:50 | 130-113-486 | Miltenyi Biotec |
| SSEA-4-VioGreen™ | REA101 | 1:50 | 130-098-341 | Miltenyi Biotec |
| SSEA5-VioBlue™ | 8e11 | 1:50 | 130-106-657 | Miltenyi Biotec |
| SOX2-FITC | REA320 | 1:50 | 130-120-721 | Miltenyi Biotec |
| OCT3/4-APC | REA622 | 1:50 | 130-123-257 | Miltenyi Biotec |
| PAX6-APC | REA507 | 1:50 | 130-123-328 | Miltenyi Biotec |
| SOX17-FITC | REA701 | 1:50 | 130-111-147 | Miltenyi Biotec |
| CXCR4-APC | REA649 | 1:50 | 130-123-814 | Miltenyi Biotec |
| CD140b-Vio®B515 | REA634 | 1:50 | 130-131-564 | Miltenyi Biotec |
| CD144-FITC | REA198 | 1:50 | 130-123-932 | Miltenyi Biotec |
| PSC Analysis Cocktail Kit, anti-human |  | 1:10 | 130-136-292 | Miltenyi Biotec |
| **Immunocytochemistry** |  |  |  |  |
| TRA-1-60-PE | REA157 | 1:25 | 130-122-921 | Miltenyi Biotec |
| OCT3/4-Vio®B515 | REA622 | 1:25 | 130-131-113 | Miltenyi Biotec |
| SOX2-Vio®R667 | REA320 | 1:25 | 130-128-344 | Miltenyi Biotec |
| CD144-APC | REA225 | 1:25 | 130-103-377 | Miltenyi Biotec |
| SM-MHC-PE | REA1107 | 1:25 | 130-119-314 | Miltenyi Biotec |
| CXCR4-APC | 12G5 | 1:25 | 130-123-814 | Miltenyi Biotec |
| SOX17-Vio®B515 | REA701 | 1:25 | 130-111-031 | Miltenyi Biotec |
| PAX6-PE | REA507 | 1:25 | 130-123-250 | Miltenyi Biotec |
| **Cyclic immunofluorescence staining (reprogramming)** |  |  |  |  |
| CD49a-PE | REA1106 | 1:50 | 130-119-306 | Miltenyi Biotec |
| TRA-1-60-PE | REA157 | 1:20 | 130-122-921 | Miltenyi Biotec |
| FSP1-PE | REA165 | 1:20 | 130-126-007 | Miltenyi Biotec |
| CD13-PE | REA236 | 1:20 | 130-120-312 | Miltenyi Biotec |
| TRA-1-81-PE | REA246 | 1:20 | 130-123-334 | Miltenyi Biotec |
| CD90-PE | REA897 | 1:50 | 130-122-864 | Miltenyi Biotec |
| FN-PE | REAL555 | 1:50 | 130-122-864 | Miltenyi Biotec |
| Ki-67-PE | REA183 | 1:50 | 130-120-417 | Miltenyi Biotec |
| NANOG-PE | REA314 | 1:50 | 130-117-377 | Miltenyi Biotec |
| wSOX2-PE | REA320 | 1:20 | 130-117-526 | Miltenyi Biotec |
| VIM-PE | REA409 | 1:50 | 130-123-774 | Miltenyi Biotec |
| OCT3/4-PE | REA622 | 1:20 | 130-120-236 | Miltenyi Biotec |

**Table S6:**
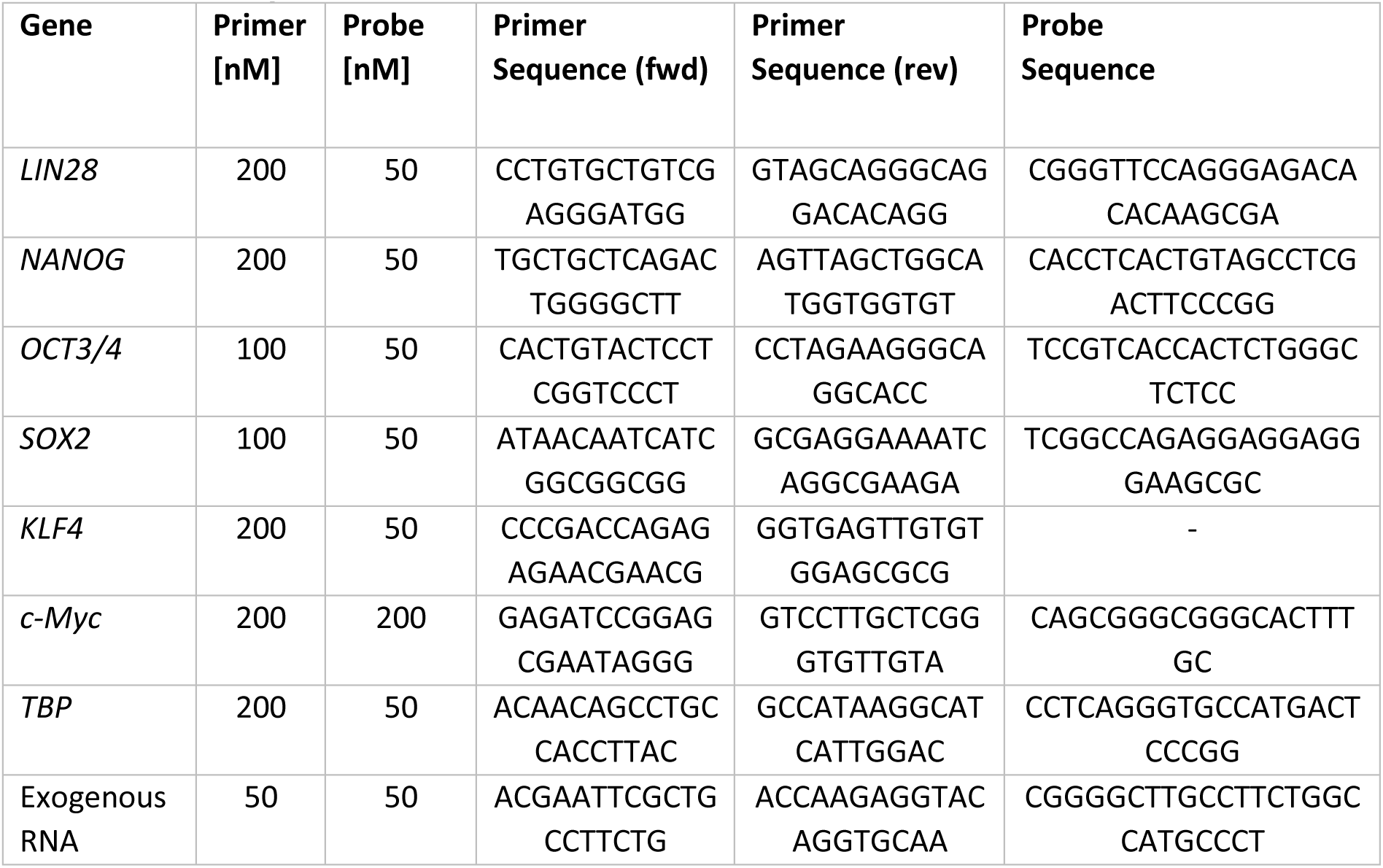
List of primers.

